# Comprehensive and Integrated Genomic Characterization of Human Immunome in Cancer

**DOI:** 10.1101/2020.06.02.128884

**Authors:** Yongsheng Li, Todd Triplett, Brandon Burgman, Ming Sun, Daniel J. McGrail, Dan Qi, Sachet Shukla, Erxi Wu, Catherine J. Wu, Anna Capasso, S. Gail Eckhardt, George Georgiou, Bo Li, Nidhi Sahni, S. Stephen Yi

## Abstract

Genetic alterations in immune-related pathways are common hallmarks of cancer. However, to realize the full potential of immunotherapy, a comprehensive understanding of immune networks and how mutations impact network structure and functional output across cancer types is instrumental. Herein we systematically interrogated somatic mutations that could express neoantigens and alter immune responses in cancer patients compared to wild-type controls. To do so, we developed a network-based immunogenomics model (NIPPER) with scoring systems to prioritize critical genes and mutations eliciting differential HLA binding affinity and alternate responses to immunotherapy. These mutations are enriched in essential protein domains and often alter tumor infiltration by immune cells, affecting T cell receptor repertoire and B cell clonal expansion. Furthermore, we devised an interactome network propagation framework integrated with drug associated gene signatures to identify potential immunomodulatory drug candidates. Together, our systems-level analysis results help interpret the heterogeneous immune responses among patients, and serve as a resource for future functional studies and targeted therapeutics.

**Significance:** Cancer cells induce specific immune-related pathway perturbations by mutations, transcriptional dysregulation, and integration of multi-omics data can help identify critical molecular determinants for effective targeted therapeutics.

## Introduction

Immunotherapy has been considered as a promising strategy for treatment of various types of cancer (1,2). Although these immunotherapy methods have made remarkable success, only a minority of patients are observed to respond to the treatment with cytotoxic T lymphocyte antigen-4 (CTLA-4) or programmed death receptor-1 (PD-1) blockade (3,4). There is a clinical need to identify predictive biomarkers and to understand the potential mechanisms of immunotherapy resistance.

With the development of high-throughput sequencing technology, recent evidence has pointed to a number of predictive biomarkers, such as tumor mutation burden (TMB) (5,6), neoantigen burden (7), PD-L1 or PD-L2 mRNA expression (8,9), epigenetic markers and immune cell infiltration profiles (10,11). With these predictive biomarkers in hand, machine learning-based methods are being built to predict the immunotherapy response. However, it is challenging to integrate different biomarkers. Moreover, increasing studies are beginning to reveal the underlying mechanisms of immunotherapy resistance. The expression of immune-related genes (11), insufficient immune cell infiltration (2), oncogenic pathway activity (12) as well as metabolism dysregulation (13) have been related to therapy resistance. But additional insights are still needed for a comprehensive understanding of the mechanisms of immunotherapy resistance.

In addition, genes do not function in isolation but rather interact extensively with each other, and the genetic abnormality is not restricted to the gene product that it encodes (14). The emerging tools of network- or pathway-based medicine offer a valuable platform to explore the predictive biomarkers as well as to understand the dysregulation pattern of cancer-related pathways. Previous studies by The Cancer Genome Atlas (TCGA) have systematically analyzed the alteration landscape of cancer-related signaling pathways (15). However, we still lack the knowledge about the alteration pattern of immune-related pathways across cancer type.

To further refine both the genomic and transcriptomic derived biomarkers for immune response, we first charted the detailed perturbation landscape of immune-related pathways. A network-based immunogenomics model with scoring systems was proposed to prioritize genes and mutations that could express neoantigens and alter immune responses in cancer patients. We identified mutations eliciting differential HLA binding affinity and these mutations are enriched in essential protein domains and often alter tumor infiltration by immune cells. Furthermore, using a network propagation framework integrated with drug associated gene signatures we discovered potential immunomodulatory drug candidates. These results can help elucidate the functionally relevant mechanisms of immune-related pathway alterations and might inform potential immunotherapy treatment options.

## Materials and Methods

### Human Immunome

Human immune-related genes were obtained from the ImmPort project (http://www.immport.org) (16). In total, 1811 genes in 17 immune-related pathways were obtained. In addition, we also obtained genes that are identified to be essential to immune response from recent literature (17,18). These genes were defined as essential immunotherapy genes and grouped into the 18^th^ pathway. In total, 2,273 immune-related genes were obtained.

### TCGA Patient Cohort

The results in our analysis are based upon datasets generated by TCGA Research Network (http://cancergenome.nih.gov/). We analyzed 33 different TCGA projects, each project represents a specific cancer type, including KIRC, kidney renal clear cell carcinoma; KIRP, kidney renal papillary cell carcinoma; KICH, kidney chromophobe; LGG, brain lower grade glioma; GBM, glioblastoma multiforme; BRCA, breast cancer; LUSC, lung squamous cell carcinoma; LUAD, lung adenocarcinoma; READ, rectum adenocarcinoma; COAD, colon adenocarcinoma; UCS, uterine carcinosarcoma; UCEC, uterine corpus endometrial carcinoma; OV, ovarian serous cystadenocarcinoma; HNSC, head and neck squamous carcinoma; THCA, thyroid carcinoma; PRAD, prostate adenocarcinoma; STAD, stomach adenocarcinoma; SKCM, skin cutaneous melanoma; BLCA, bladder urothelial carcinoma; LIHC, liver hepatocellular carcinoma; CESC, cervical squamous cell carcinoma and endocervical adenocarcinoma; ACC, adrenocortical carcinoma; PCPG, pheochromocytoma and paraganglioma; SARC, sarcoma; LAML, acute myeloid leukemia; PAAD, pancreatic adenocarcinoma; ESCA, esophageal carcinoma; TGCT, testicular germ cell tumors; THYM, thymoma; MESO, mesothelioma; UVM, uveal melanoma; DLBC, lymphoid neoplasm diffuse large B-cell lymphoma; CHOL, cholangiocarcinoma.

### Molecular and Clinical Datasets across Cancer Types

Somatic mutations were obtained from the publicly available TCGA MAF file (“MC3”) which covers 10,224 patients (19). This dataset was directly downloaded from Synapse under the number of syn7214402. Six calling methods were applied and numbers of filters were applied. All these mutations were subjected to ANNOVAR(20) and we obtained the conservation and MetaSVM score for each mutation (21).

RNA-seq data were obtained from the TCGA project via the R-package “TCGAbiolinks” (22), which was specifically developed for integrative analysis with GDC data. We downloaded the Fragments Per Kilobase of transcript per Million mapped reads (FPKM)-based gene expression for 33 types of cancer. In addition, we also obtained the raw read counts for each gene in these samples.

The clinical information for patients of 33 cancer types were downloaded from TCGA project via the R-package “TCGAbiolinks”, including the survival status, stages, grades, survival time.

### Tumor Purity Estimation and Immune Cell Deconvolution

ESTIMATE (Estimation of STromal and Immune cells in MAlignant Tumor tissues using Expression data) was used to estimate the immune score, stromal score and tumor purity of each patients across 33 cancer types (23). The FPKM-based gene expression profiles were used as input of this method. In addition, TIMER (https://cistrome.shinyapps.io/timer/) was used to estimate the abundances of member cell types in a mixed cell population, using gene expression data (7).

### Pathway Mutation Burden Score

Pathway Mutation Burden (PMB) score was defined to evaluate whether the immune pathways were likely to mutate in a specific cancer type. This score considers the proportion of mutated genes as well as the coverage of patients in each cancer type. For an immune pathway *i* in cancer type *j*, the score is defined as following:

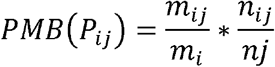

Where *m*_*ij*_ is the number of mutated immune genes in pathway *i*; *m*_*i*_ is the total number of genes in pathway *i*; *n*_*ij*_ is the number of patients of cancer *j* that with gene mutations in pathway *i*; and *n*_*j*_ is the total number of patients sequenced in cancer *j*.

### Immune Response-related Scoring System

Here, we considered three immune response-related scores estimated from gene expression. The immune score that represented the infiltration of immune cells in tumor tissue was assessed based on ESTIMATE (23). The MHC score was formulated from the gene expression levels of the “core” MHC-I set (including HLA-A, HLA-B, HLA-C, TAP1, TAP2, NLRC5, PSMB9, PSMB8 and B2M) (24). The FPKMs of genes were first log-transformed and then median-centered. The mean expression levels of these core genes was defined as the MHC score. The CYT score was a quantitative measure of immune cytolytic activity based on transcript levels of two key cytolytic effectors, granzyme A (GZMA) and perforin (PRF1), found in previous studies (25).

### Classification of Patients with Different Immune Responses

To investigate whether the immune-related scores could be used to classify the patients with different immune responses to immunotherapy, we obtained gene expression profiles of patients treated with MAGE⍰A3 immunotherapy in metastatic melanoma patients (25). The three types of immune response-related scores were calculated for each patient. Next, we used the Receiver Operating Characteristic (ROC) Curve Analysis to evaluate the effects of these scores. This process was performed by using ‘ROCR’ package in R program.

### Identification of Protein Regions Associated with Immune Response

The domainXplorer algorithm was used to identify the protein regions that are associated with immune scores (26). The protein functional regions were defined as protein domains in Pfam (27) or intrinsically disordered regions identified by Foldindex (28). In addition, potential domains identified by AIDA (29) were also included in our analysis. Here, we only focused on immune genes-related functional regions.

In brief, domainXplorer used three statistical tests to evaluate whether mutations in a functional protein region were correlated with the immune response-related scores (26). The first linear model was defined as follow:

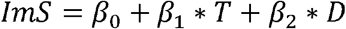

Where “*ImS*” is the immune score of each patient, “*T*” is the tissue of origin of each patient (here is the cancer type), and “*D*” indicates whether the functional region is mutated or not. In the next step, Wilcox rank sum test was used to compare the immune response-related scores in patients with mutations in the functional region being analyzed against those with mutations in other regions of the same immune gene (Test-2) or those without mutations in the immune gene at all (Test-3). Protein regions with p1<0.05, p2<0.05 and p3<0.05 were identified as to be associated with immune response. In addition, we also identified the protein regions that were correlated with the proportion of immune cell infiltration based on the same method.

### Differential Expression Analysis

To identify differentially expressed genes in each cancer type, we downloaded raw RNA-seq counts for all tumor samples. We then used limma (30) to identify the genes and considered the tumor purity as a factor. Genes with p-values less than 0.05 and four-fold changes in expression were identified as differentially expressed genes in each cancer type.

To evaluate the similarity of cancer types based on the differentially expressed immune genes, we calculated the Jaccard index for each pair of cancer

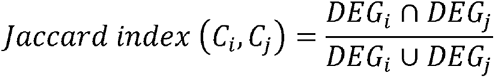

Where *DEG*_*i*_ and *DEG*_*i*_ are the differentially expressed immune genes in cancer type *i* and *j*.

### Gene Set Enrichment Analysis

Gene Set Enrichment Analysis was used to identify the enriched immune pathways by differentially expressed genes in each cancer type (31). First, genes were ranked by negative log10 of the differential expression analysis-derived FDR multiplied by the sign of the logFC (log fold change). The “weighted” enrichment statistics were calculated for enrichment or depletion of each immune pathway in specific cancer type.

### Prediction of Neoantigens and Functional Screen of Mutations

The somatic mutations were collected from TCGA project and we obtained the HLA alleles for tumor samples from TCIA (https://tcia.at/home) (32). Next, NetMHCpan was employed for neoantigen prediction based on the information on mutations and HLA alleles (33). All the wild-type and mutated peptides with 8-11 amino acids were extracted and predicted for the binding affinity. The strong binding is defined as IC50<150 nM and weak binding with 150 nM<IC50<500 nM based on the previous study (34).

### Network-based Prioritization of Immunome-related Genes and Drug Discovery

Human protein-protein network was integrated into our analysis to prioritize immune dysregulated genes. First, we extracted the subnetwork formed by immune-related genes from an integrated functional network (35). The main component of this subnetwork consists 6,060 interactions among 1,202 immune-related genes. Next, the immune genes were ranked based on the correlation between gene expression and immune scores, MHC scores and CYT scores. Top 50 genes with higher positive- or negative-correlation with immune, MHC and CYT scores were selected as seed genes. In addition, the genes which mutations are associated with immune-related scores were also integrated as seed genes. To simulate the propagation of the genetic alterations through the network, we employed the random walk with restart (36). When an immune gene in the network is ranked in top 50, a value “1” is initially assigned to the gene. Then the genetic alterations are propagated along the neighbors such that the genetic alteration probability was calculated for all genes in the network according to the following equation:

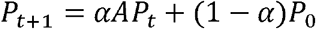

Where *P*_0_ is the initial binary probability vector, A denotes a degree-normalized adjacency matrix of the immune subnetwork, and *α* determines the diffusion degree of the genetic influence through the network. We used the optimal value (*α* = 0.7) in our analysis (37). After numbers of iteration, we obtained the final probability for each gene in the network. Genes with greater score than average probability were identified as immune dysregulated genes for each cancer type (38).

Next, we integrated drug associated gene signatures to identify the drug candidates for each cancer type. We first obtained the expression signature genes from The Connectivity Map (also known as CMAP) (39). For each drug, there are two types of signature genes, one is the up-regulated genes and the other one is the down-regulated genes. We used hypergeometric test to evaluate whether the drug-perturbed gene signatures were enriched in cancer-specific immune gene sets. For each cancer type, we obtained the p-value and then all the p-values across 33 cancer types were combined to a combined p-value.

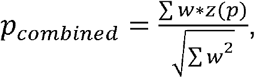

Where w is the weight for the individual p-values, and z is the z-score of p-values. Here, we set the same weights for all individual p-values. The drugs with adjusted p-values less than 0.01 were identified. We obtained the drugs that had up-regulated/down-regulated gene signatures enriched for genes positively/negatively correlated with immune scores.

### Survival Analysis

The patients were divided into two groups based on the median expression of specific immune gene. Log-rank test was used to evaluate the survival difference between two groups. Genes with p-values less than 0.05 were considered as significant.

## Results

### Genome and Transcriptome-based Immune Response Prediction

Tumor mutational burden (TMB), a measurement of the overall number of genetic alterations observed in a cancer sample, has been suggested as a potentially prognostic marker for immunotherapy (40,41). In addition, several gene signatures have also been proposed to predict the immune response of patients (Figure 1A). We first evaluated whether the three immune-related scores (immune score, MHC score and cytolytic activity) could be used to predict the immune response of tumor patients. Based on the gene expression data in response to treatment with MAGEl7l A3 immunotherapy of metastatic melanoma patients (42), we found that the patients that responded to immunotherapy had significantly higher immune scores (p=0.003), MHC scores (p=1.7e-4) and cytolytic activity (p=0.007, Figure 1B). Moreover, we found that these three immune-related scores indeed distinguished the responder from non-responder patients with area under the curve of the receiver operating characteristic (AUROC) from 0.71 to 0.79 (Figure 1C). To further validate this, we analyzed another gene expression dataset for PD-1 immune checkpoint blockade (43). We also found that patients which were responsive to the PD-1 immunotherapy exhibited higher immune-related scores (Figure S1A). The AUROCs for classification ranged from 0.52 to 0.85 (Figure S1B), which was similar to the results obtained from gene signatures in previous studies (44). These results suggest that the immune-related scores can potentially reflect the immune milieu of tumors in patients and help identify specific factors that may modulate these environments.

**Figure-1.**
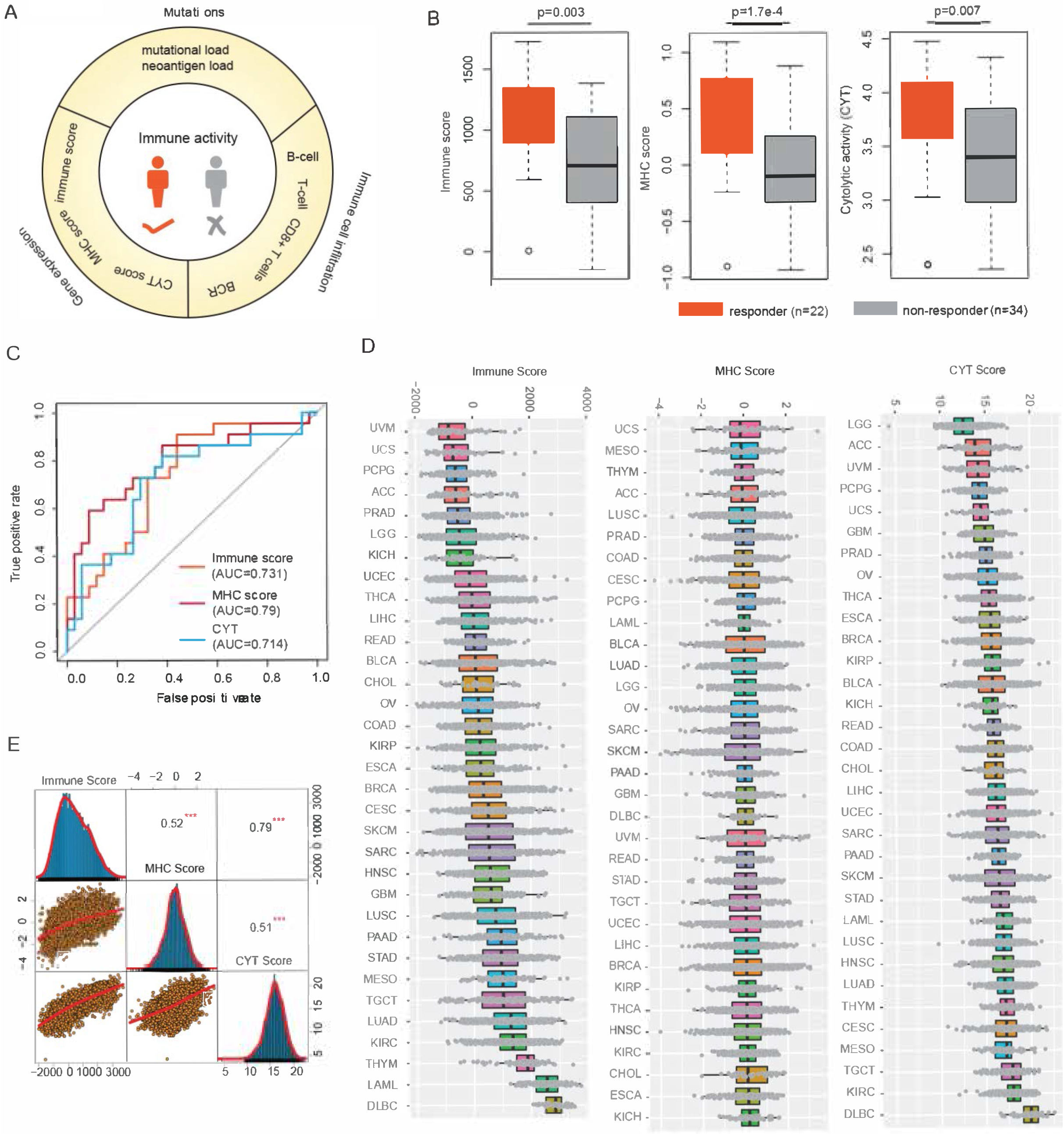
Immune response prediction signatures in cancer. (A) Literature curated immune response prediction signatures. (B) Boxplots show the distribution of Immune score, MHC score and CYT score in immune response vs non-response patients. (C) Receiver operating characteristics (ROC) analysis of immune score, MHC score and CYT score from each prediction. (D) The distribution of immune score, MHC score and CYT score across 33 cancer types in TCGA cohort. Cancers are ranked by the median scores. (E) Scatter plots show the correlation between, immune score, MHC score and CYT score across cancer types. ***p<0.001.

We next explored the generalizability of these scoring systems in predicting patient immune responses across cancer types. We first calculated the three immune-related scores (immune score, MHC score and cytolytic activity) for all tumor patients from the TCGA project (Figure 1D). We found that the distributions of the immune-related scores were diverse across 33 cancer types. Specifically, patients in several cancer types, such as lymphoid neoplasm diffuse large B-cell lymphoma (DLBC) and kidney renal clear cell carcinoma (KIRC), exhibited higher immune-related scores. These results suggest that immunotherapy might be suitable for these cancer types. Moreover, although the three immune-related scores appeared to be correlated with each other, the Pearson Correlation Coefficients (PCC) ranged from 0.51-0.79 (Figure 1E). This observation implied that these scores might be complementary with each other, therefore integrating these scores would facilitate the identification of novel candidate gene targets for immunotherapy.

### Immune Mutational Burden Analysis Identifies Critical Pathways and Genes

It has been suggested that the mutations in immune pathways could impact tumor-immune interactions (45,46). It is unclear, however, how the immune mutational burden (IMB) is associated with patient outcome. Moreover, it seems that the mutation burdens across pathways are varied (15). Therefore it remains elusive if and how mutations in immune-related pathways contribute to higher mutation burden. To systematically investigate the global functional genomic landscape of immune-related genes, we manually curated 2,273 immune-related genes from literature (16,17). These genes were further classified into 18 immune-related pathways (Figure 2A). Next, we analyzed whole exomes from 100 melanoma patients treated with CTLA-4 blockade (47). We first calculated the number of mutations (tumor mutation burden, TMB) and the number of mutations located in immune-related genes (defined as immune mutation burden, IMB). We further compared TBM and IMB between responders and non-responders. The mutation burden and immune burden of responders were significantly higher than those of non-responders (Figure S2A, p<0.05). In addition, mutation burden and immune burden could predict immune response with similar power (Figure S2B, AUCs=0.68). Furthermore, we calculated the IMB for each patient in TCGA cohort. We found that the cancer types that are likely to respond well to immunotherapy had higher proportion of immune-related gene mutations, such as skin cutaneous melanoma (SKCM) and kidney cancer (Figure S2C). These results suggest that the immune burden is a useful indicator of immune response; target sequencing of immune-related genes would help predict the immune response of patients.

**Figure-2.**
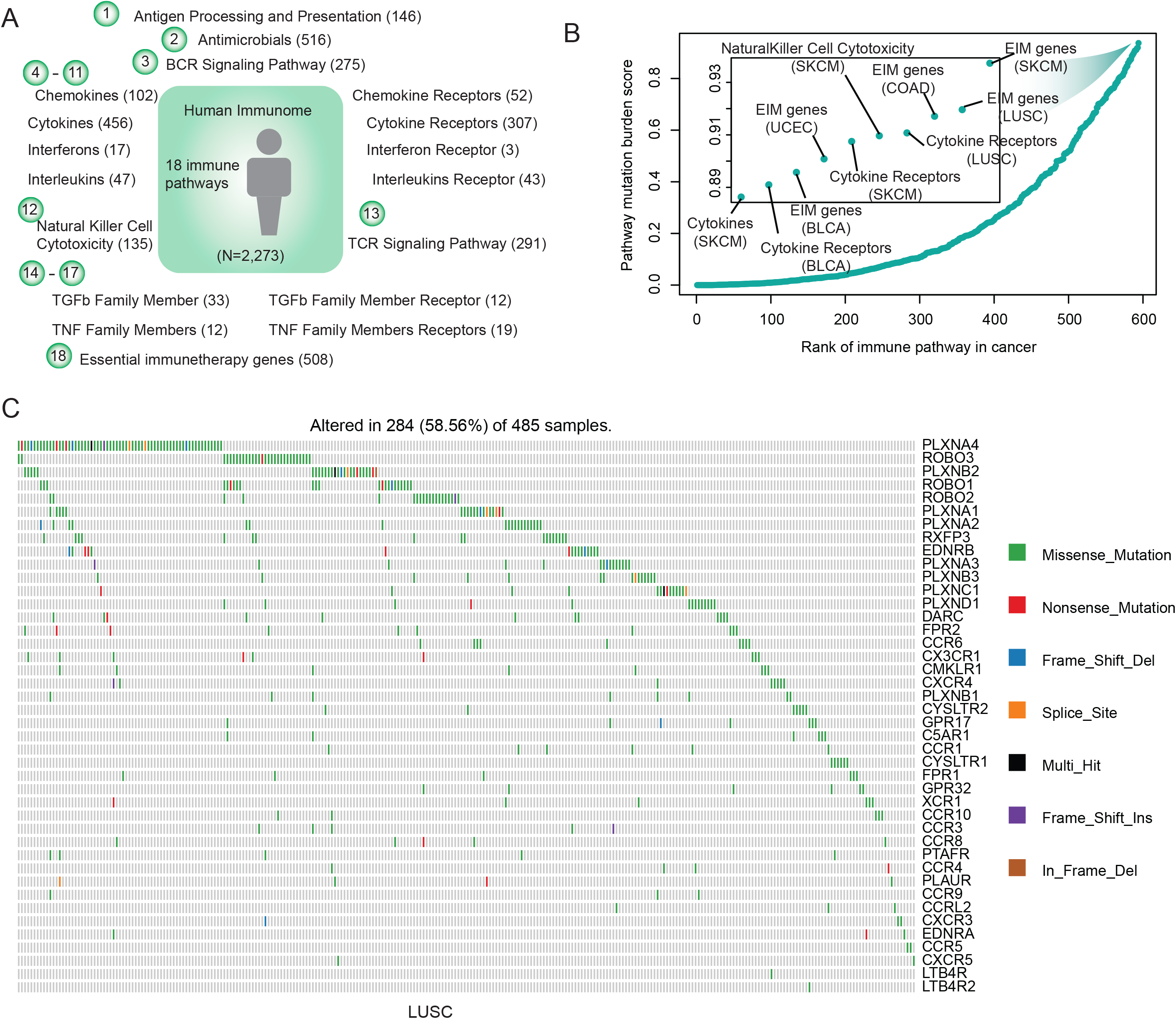
Critical mutated immune pathway and genes in cancer. (A) Human immune pathways and number of genes in each pathway. In total, 2,273 immune-related genes in 18 pathways are analyzed. (B) The PMB score for each immune-related pathway in cancer. Each dot represents a pathway in a specific cancer type. The immune score is calculated as the product of the proportion of mutated genes in the pathway and the percentage of mutated samples. Pathways are rank by the PMB score. (C) The mutations of genes in cytokine receptor pathway in LUSC. Genes were ranked by the mutation frequency.

Next, we defined a Pathway Mutation Burden (PMB) score to evaluate to what extent each immune-related pathway is perturbed by genomic alterations. This score is calculated based on the mutation frequency and the proportion of mutated genes in each pathway. We found that the essential genes identified by CRISPR-Cas9 KO screens (17) had higher PMB scores across cancer types (Figure 2B). Moreover, we observed another pathway-cytokine receptors (CRs) with higher PMB scores in LUSC, SKCM and DLBC (Figure 2B). We next focused on the CR mutations in LUSC (Figure 2C). Consistent with previous studies, the mutations in CR pathway exhibited a mutual exclusive pattern (48,49). Moreover, the mutations in CR pathway had higher functional impact scores than other mutations evaluated by metaSVM (21) (Figure S2D). The top-1 gene (PLXNA4) ranked by mutation frequency was mutated in approximately 19% of lung cancer patients. PLXNA4 was shown previously to promote tumor progression and tumor angiogenesis by activating VEGF and FGF signaling (50). We further found that the expression of PLXNA4 was correlated with the patients’ survival in LUSC (Figure S2E, p=0.017). These results suggest that the mutations in cytokine receptors pathway might play critical roles in cancer and the proposed PMB index could help prioritizing key mutations in cancer pathways.

### Widespread Transcriptional Alterations of Human Immunome across Cancer Types

Next, we interrogated if and to what extent the human immunome is altered in cancer at the gene expression level. We identified differentially expressed genes across 18 cancer types with more than five normal samples. As tumor purity may affect the identification of differentially expressed genes (51), we first estimated the tumor purity of each sample (Figure S3A) and identified the differentially expressed genes using tumor purity as a co-variable in the regression model. We found that the immune-related genes were significantly perturbed in cancer when compared to other coding genes (Figure 3A and Figure S3B, p-values<0.05 for all cancer types, Fisher’s exact test). The Odd Ratios (ORs) of immune genes vs non-immune genes were higher than 1.0 in all cancer types. These results indicate that these genes could play a role in promoting tumor growth.

**Figure-3.**
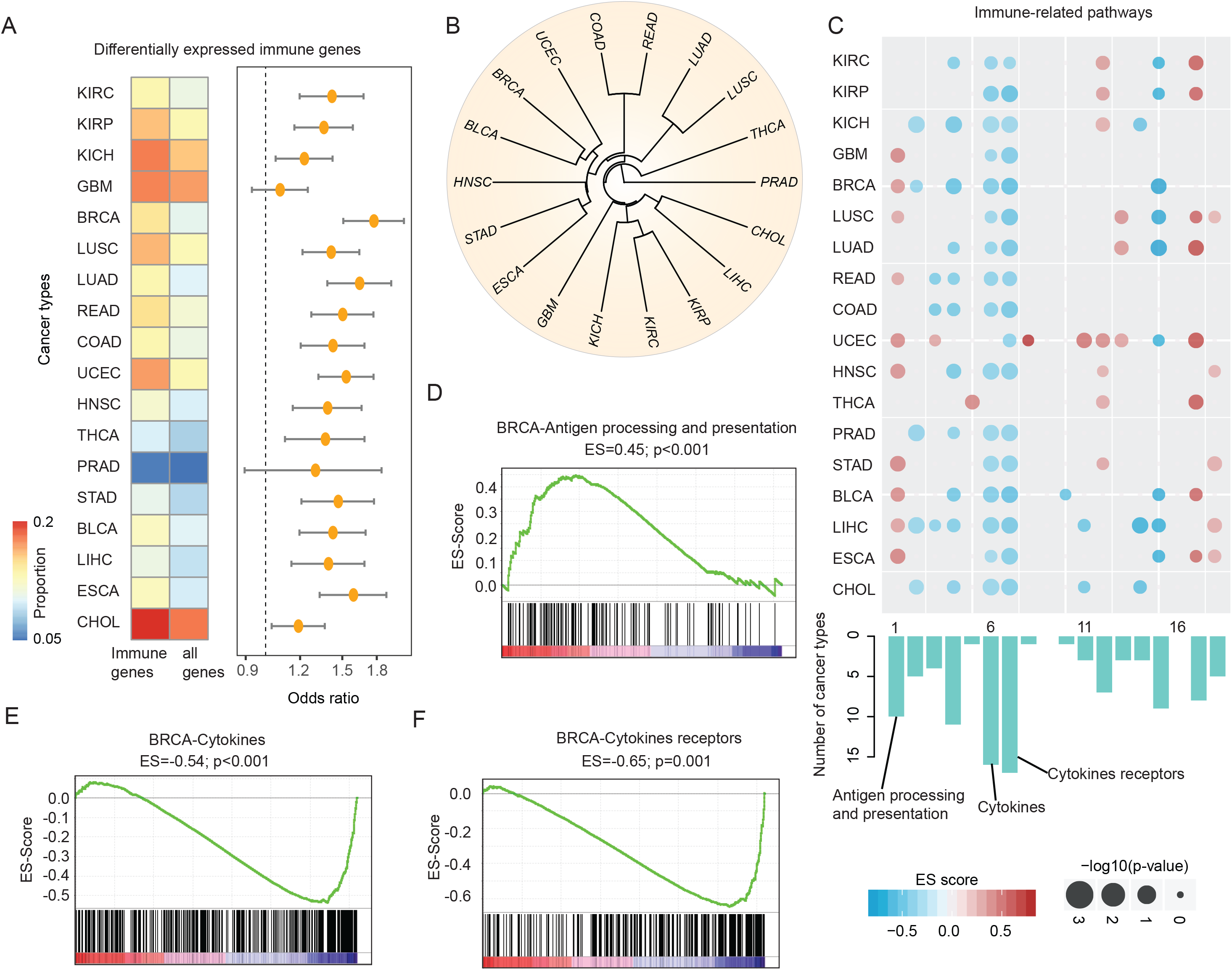
Widespread transcriptome alterations of human immunome in cancer. (A) The proportion of differentially expressed immune genes and all human genes. The odds ratios and 95% confidence levels of Fisher’s exact tests are shown in the right panel. All p-values are less than 0.05 for 18 cancer types with at least fiver normal samples. (B) Cluster of cancer types based on the similarity (Jaccard index) of differentially expressed immune genes. Cancer types of similar tissue origin are cluster together. (C) Heat map shows the p-values and enrichment score for the differentially expressed genes enriched in immune-related pathways. Each row represents a cancer type and each column represents an immune-related pathway. The sizes of dots show the - log (p-values) and the color of dots show the enrichment score. The bar plots below the heat map show the number of cancer types enriched by each pathway. (D) - (F) Gene Set Enrichment Analysis shows enrichment repression of cytokines, cytokines receptors and activated antigen processing and presentation in breast cancer. The horizontal bar in graded color from red to blue represents the rank-ordered, non-redundant list of all genes. The vertical black lines represent the projection of immune genes in each pathway onto the ranked gene list. (D) for antigen processing and presentation pathway, (E) for cytokines pathway and (F) for pathway cytokines receptors in breast cancer.

An increasing number of studies have demonstrated that cancer types with similar tissue origin show similar transcriptome and regulation (52,53). Next, we calculated the Jaccard index for each pair of cancer based on the similarity of differentially expressed immune-related genes (see details in Methods). We found that cancer lineages with similar tissue origin were clustered together (Figure 3B), such as lung adenocarcinoma (LUAD) and lung squamous cell carcinoma (LUSC), cholangiocarcinoma (CHOL) and liver hepatocellular carcinoma (LIHC), and three types of kidney cancer. These results indicate that these cancer types show similar immunome perturbations.

In addition, we ranked all genes based on their differential expression and used gene set enrichment analysis (GSEA) to explore which immune-related pathways were likely to show transcriptional perturbation (31,54). 16/18 immune-related pathways were significantly altered in at least one cancer type (Figure 3C). The antigen processing and presentation, cytokines and cytokines receptors pathways were perturbed in more than 16 cancer types. Moreover, we found that these immune-related pathways showed consistent trend (enriched or depleted) across cancer types. For instance, antigen processing and presentation pathway was likely to be activated in breast invasive carcinoma and glioblastoma (BRCA and GBM, Figure 3C and Figure S3C). In addition, cytokines and cytokine receptors pathways were repressed in BRCA and GBM (Figure 3E-F and Figure S3D, FDR<0.001).

### Critical Regions of Immune Genes Associated with Immune Response

Growing evidence suggests that mutations in certain genes influence the immune response (26). Identification of these genes may provide valuable insights for improving immunotherapy. Thus, we explored ‘domainXplorer’ to identify critical regions of immune genes that are associated with immune response in cancer (26). Given the importance of the proposed immune-related scores in predicting immune response, we next determined whether cancer mutations in immunity candidate driver regions were significantly correlated with these scores. Our analysis yielded a total of 209 protein regions in 145 immune-related genes that are potential cancer immunity drivers (Figure 4A). Specifically, mutations in seven genes (UBXN1, LTBP2, STAT1, NGFR, NRAS, MET and LTBP4) were significantly correlated with all of three types of immune-related scores. LTBP4, a member of the latent TGFβ binding protein gene family, had been linked to several human diseases (55). We found that patients with EGF_CA domain mutations exhibited significantly higher immune scores, MHC scores and cytolytic activity (Figure 4B, p<0.05). Moreover, we also identified mutations of STAT1 significantly correlated with these scores, which was consistent with previous observation that STAT1 played an important role in the innate immune response (56). These results collectively expand the catalog of potential cancer immune drivers and highlight the importance of taking into account the protein structural context to identify patients that are likely to clinically benefit from immunotherapy.

**Figure-4.**
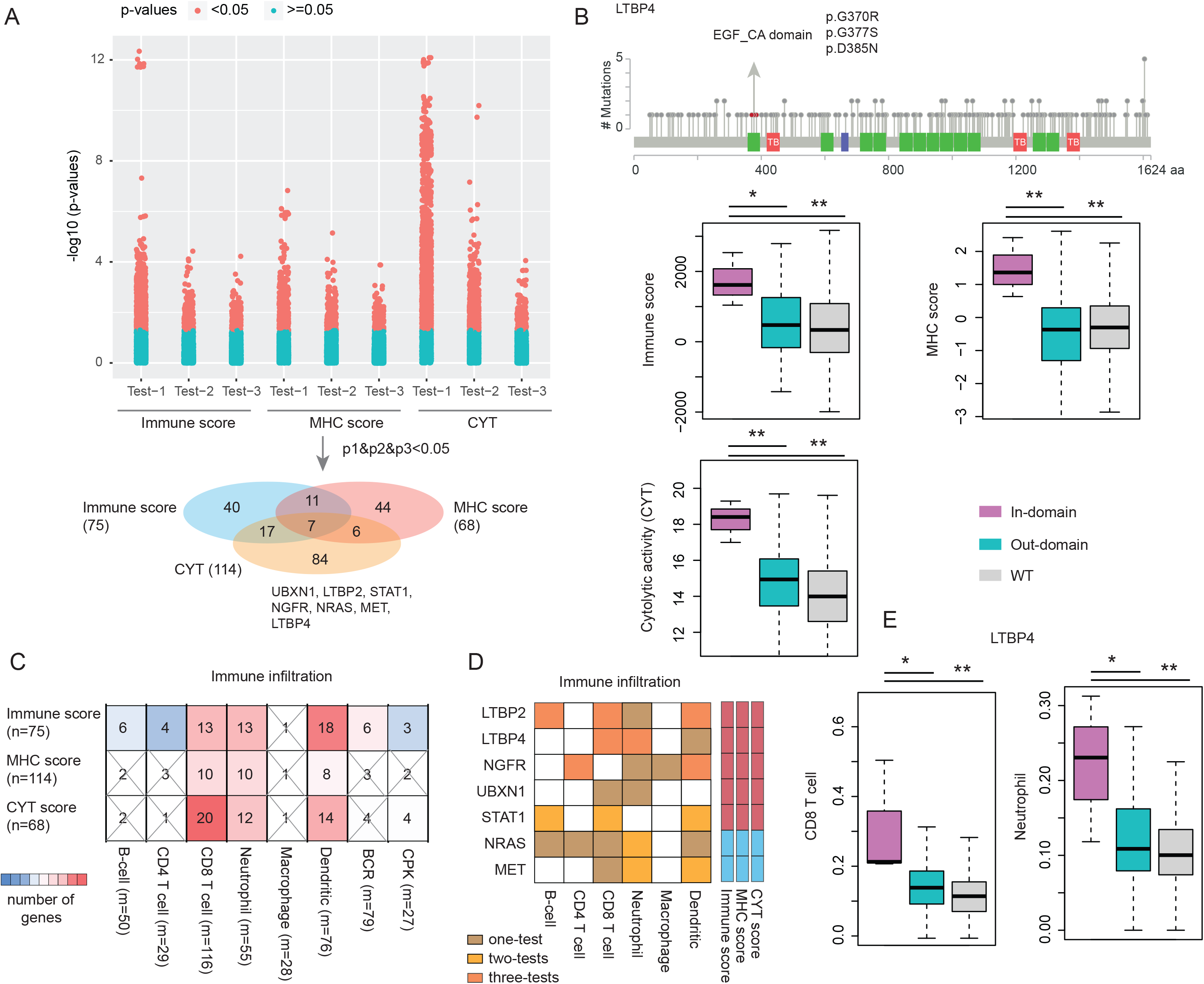
Critical protein-regions related to immune response in cancer. (A) The distribution of p-values for testing the association of protein regions and immune response scores. The bottom Venn plot shows the overlap of protein-regions that are related with immune score, MHC score and CYT score. The seven overlap genes were shown. (B) Representative mutated protein regions that are related with immune response in cancer. The mutations in the EGF_CA domain of LTBP4 gene are shown. The up-panel shows the mutations in the LTBP4 and the domain region is marked by green and mutations are shown as red balls. The immune score, MHC score and CYT score distributions of the samples with domain mutations, other region mutations and wild-type are shown in the bottom-panel. ***, p<0.01, Wilcox’s rank sum test. (C) The overlap of protein regions which mutations were related with immune response scores and immune cell infiltration. The rectangle colored are with p<0.05. (D) The overlapped seven genes mutation associated with distinct immune cell infiltration. (E) The distribution of immune cell infiltration for patients with in-domain mutations, out-domain mutations and wild type of LTBP4 gene. Left for CD8 T cell and right for Neutrophil cell.

### Somatic Mutations Associated with Immune Cell Infiltration

Our above analyses identified candidate immune driver protein regions that are potentially correlated with immune response. However, we are still lack of knowledge about the potential mechanism. An emerging role in treatment response has been attributed to immune cell infiltration in human tumors (57). Thus, we next explored to what extent the mutations in these regions were related with immune cell infiltration. We first identified the protein regions whose mutations were significantly correlated with the abundance of six tumor-infiltrating immune cells (TIIC) subsets (B cells, CD4 T cells, CD8 T cells, macrophages, neutrophils, and dendritic cells) (58). We also identified the proteins that were associated with B cell receptor (BCR) diversity and CPK (TCR/BCR CDR3s per kb of TCR/BCR reads) used to evaluate the clonotype diversity of T and B cells (59). We found that both MHC and CYT scores showed significant association with the expansion of tumor-infiltrating B cells after correction for tumor purity (Figure S4A). In addition, MHC score was weakly associated with TCR diversity, potentially due to clonal expansion of tumor antigen-specific T cell populations. CYT score on the other hand was strongly correlated with CPK (Figure S4B). We found that the protein regions that were correlated with immune score, MHC score and cytolytic activity were significantly overlapped with those correlated with TIIC abundance (Figure 4C). Especially, there were a higher number of protein regions correlated with CD8 T cells, neutrophils, and dendritic cells.

Specifically, we found that seven genes, whose mutations correlated with all three types of immune-related scores, were associated with at least one type of immune cell abundance (Figure 4D). For example, patients with LTBP4 mutations exhibited significantly higher abundance of CD8 T cells and neutrophils (Figure 4E). CD8+ T cells are crucial mediators of anti-tumor immune response and the targets of checkpoint blockade (60), while neutrophils are part of the innate immune system whose roles in cancer have been mixed (61). Moreover, we found that not only the genomic variants in LTBP4 were correlated with immune cell abundance, but patients with copy number alterations also exhibited significantly different abundance (Figure S4C). These results suggest that the genomic alterations might contribute to immunotherapy response through affecting the immune cell infiltration in cancer.

### Critical Immune Genes Enriched for Neoantigens in Cancer

Increasing studies have demonstrated that mutation-derived neoantigens form a major ‘active ingredient’ of successful cancer immunotherapies (62,63). Next, we predicted the neoantigens that were derived from mutations of TCGA project based on NetMHC (64). We found that the immune-related genes tended to generate significantly more neoantigens compared with coding genes in general (Figure 5A, p=1.18e-4, Fisher’s exact test). Especially we found that the mutations in PIK3CA, KRAS, HRAS and NRAS were likely to generate neoantigens. Approximate 25% of the mutations in PIK3CA and HRAS could generate neoantigens (Figure 5B). This result is consistent with the observation that RAS plays a role in immunotherapy (65,66).

**Figure-5.**
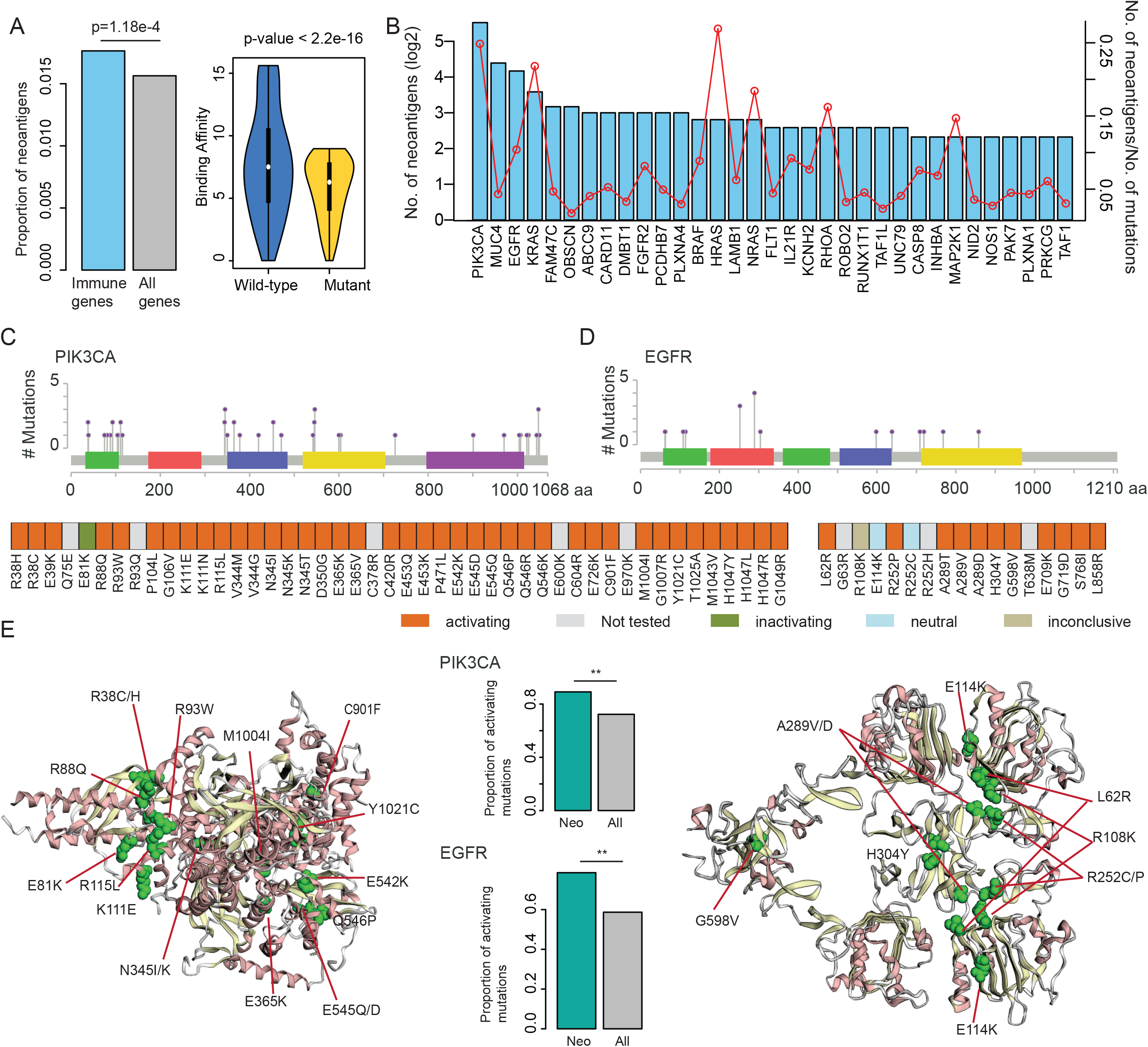
Neoantigen enriched in immune-related genes. (A) The left panel shows the proportion of neoantigens in immune gene compared to all coding genes (p=1.18e-4, Fisher’s exact test). The right panel shows the distribution of binding affinity of HLAs in wild-type and mutated peptides (p<2.2e-16, Wilcox’s rank sum test). (B) Number of neoantigens in immune-related genes. The barplots show the number of neoantigens generated from each immune-related genes. The red dot line shows the number of neoantigens/number of total mutations in each gene. (C) The mutations distribution in the PIK3CA that generate neoantigens. (D) The mutations distribution in the EGFR that generate neoantigens. (E) The left and right panels shows the 3D structure of PIK3CA and EGFR, red dots represent the mutations in the same cluster. The middle panels show the proportion of functional mutations, the green bars for mutations generated neoantigens and grey bars for all screened mutations. **p<0.05 for Fisher’s exact test.

Next, we specifically focused on mutations in PIK3CA and EGFR that give rise to neo-antigens. Based on the viability-based functional screen in two cell lines (Ba/F3 and MCF10A) (67), we found that the mutations that encode neoantigens were likely to be activating mutations in both cell lines (Figure 5C-5D). For example, we identified 46 missense mutations that could generate neoantigens in PIK3CA. Among these mutations, 41 mutations were in the functional dataset. We found that 40 mutations were activating mutations and one was inactivating mutations. Moreover, we identified a mutation cluster (including R38C/H, E81K, R88Q, R93W, K111E and R115L) in PIK3CA (Figure 5E). For EGFR, we identified 17 mutations that could generate neoantigens. In total, 14 mutations were screen for function and 11 mutations were activating, two were neutral and 1 inconclusive in two cell lines (Figure 5D). We also found seven mutations formed a cluster in 3D (including R108K, R252/P/C/H and A189V/D, Figure 5E). These results suggest that these mutations may play a role in the growth of tumors and are less likely to be selected during immune editing and undergo antigen escape.

### Prioritization of Candidate Driver Genes Based on Immunome Network

Our above results indicate widespread genomic and transcriptome perturbations in the cancer immunome network. An integrated landscape of these perturbations offers a basis for further advancing our understanding of the immune regulation in cancer. Thus, we proposed a Network-based Integrative model to <u>P</u>rioritize the <u>P</u>otential immune driv<u>ER</u>s (NIPPER) (Figure 6A). Briefly, we first identified the genes whose mutations or expression were significantly correlated with immune scores, MHC scores and CYT scores. These genes were then mapped to an immune-related protein-protein interaction (PPI) network, which consisted of 6,060 interactions among 1,202 immune-related genes. Next, the mutational effects were propagated in the PPI network. Finally, the genes with driver probability greater than expected were identified as immune response-related signature genes. This process was repeated for all cancer types. Here, the genes positively/negatively correlated with immune-related scores were analyzed, separately.

**Figure-6.**
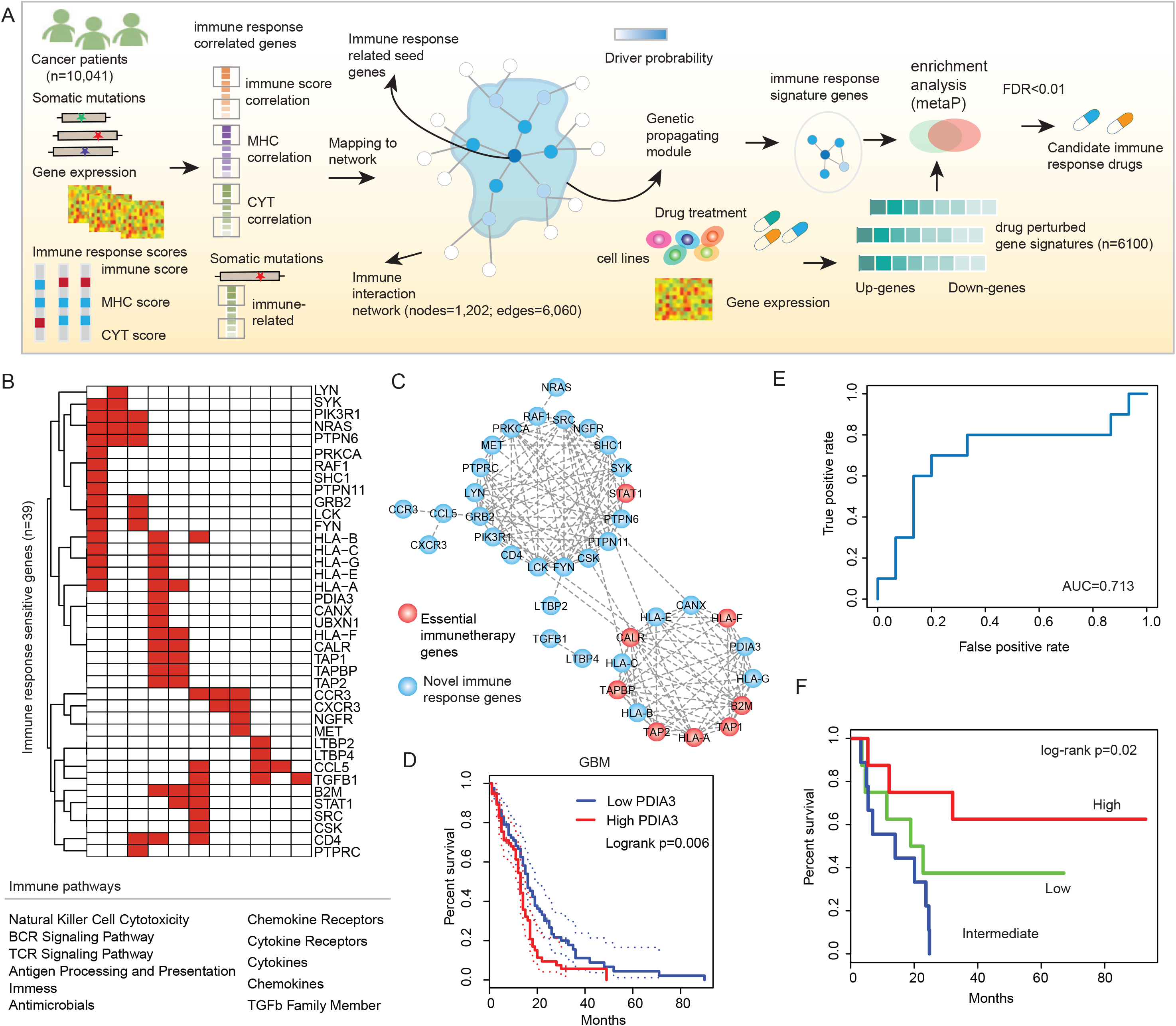
Network-based immune candidate driver identification. (A) Flowchart to identify the candidate personalized immune driver genes and the candidate drugs for each patient. For each cancer type, expression or mutations of genes that are correlated with immune response scores are mapped to an immune gene-gene interaction network. The immune response correlation is propagated via network links and immune genes are identified based on random walk method. Drug perturbed gene signatures (n=6100) are integrated for identification of the drugs that enriched the candidate immune driver genes (FDR<0.01) of the specific patient. (B) Top ranked immune response score-related genes across cancer types. Genes are ranked by the number of cancer types that show correlation between gene expression and immune response scores. (C) Top-ranked 39 genes are identified and the corresponding pathways were shown. The subnetworks formed by these 39 genes are shown in the up-panel with node colors indicate whether they are identified by the CRISPR-Cas9. (D) The expression of PDIA3 (linked more known essential immune genes) is correlated with patients’ survival in GBM. (E) ROC curve for the classification of patients response to immunotherapy based on candidate genes. (F) Kaplan-Meier survival curves for patients classified by the average expression of candidate genes.

Based on the proposed network-based model, we identified 39 genes as signature genes in 33 cancer types (Figure 6B). Patient with high expression of these genes were likely to response to immunotherapy. These genes were involved in 11 immune-related pathways, and 8 genes (such as STAT1, B2M, TAP1 and TAP2) had been identified as essential genes in immune response. Moreover, we found that these candidate immune response-related genes were clustered together as modules in the PPI network (Figure 6C). Seven of the 8 essential immune genes were linked in a dense module, and STAT1 was clustered together with more novel immune response-related genes; many of these were enzymes that play critical roles in modulating signaling by the T cell receptor (e.g. LCK, FYN, PTPN11). Interestingly, we identified one candidate gene-PDIA3, which linked 7 known essential genes. The expression of this gene was correlated with the patients’ survival in several cancer types, including CESC, LUAD, HNSC and GBM (Figure 6D and Figure S5). These observations suggest that this gene may play critical roles in immunotherapy, which is consistent with the conclusion of one recent study (68).

Next, we validated these genes in another cohort that underwent treatment with immunotherapy (24). We found that the expression of these genes could classify the responder and non-responder patients with an AUC of 0.713 (Figure 6E). Moreover, the average expression of these genes were significantly associated with improved overall survival (Figure 6F, log-rank p=0.02).

### Mutation-Perturbed HLA Binding and Potential Immune-related Drugs

Next, we evaluated mutations in the genes identified by NIPPER in different cancers. We found that the mutations in these genes were likely to decrease the binding affinity of HLAs (Figure 7A, p<0.001, Wilcox’s rank sum test). Specifically, we identified 165 mutations in 30 genes that induced the weaker binding of HLAs (Figure 7B). These mutations were distributed in various types of cancer. We identified six mutations in PIK3R1 (encoding p85α) that caused the weak binding of HLAs (Figure 7B and 7D). It had been demonstrated that both loss-of-function and gain-of-function mutations in this gene lead to immunosuppression (70). Consistently, we identified two mutations (N564D and K567E) that increased growth of cancer cell lines (Figure 7D). These results suggest that our network-based method helps identify candidate driver mutations by inducing the decreased binding of HLAs.

**Figure-7.**
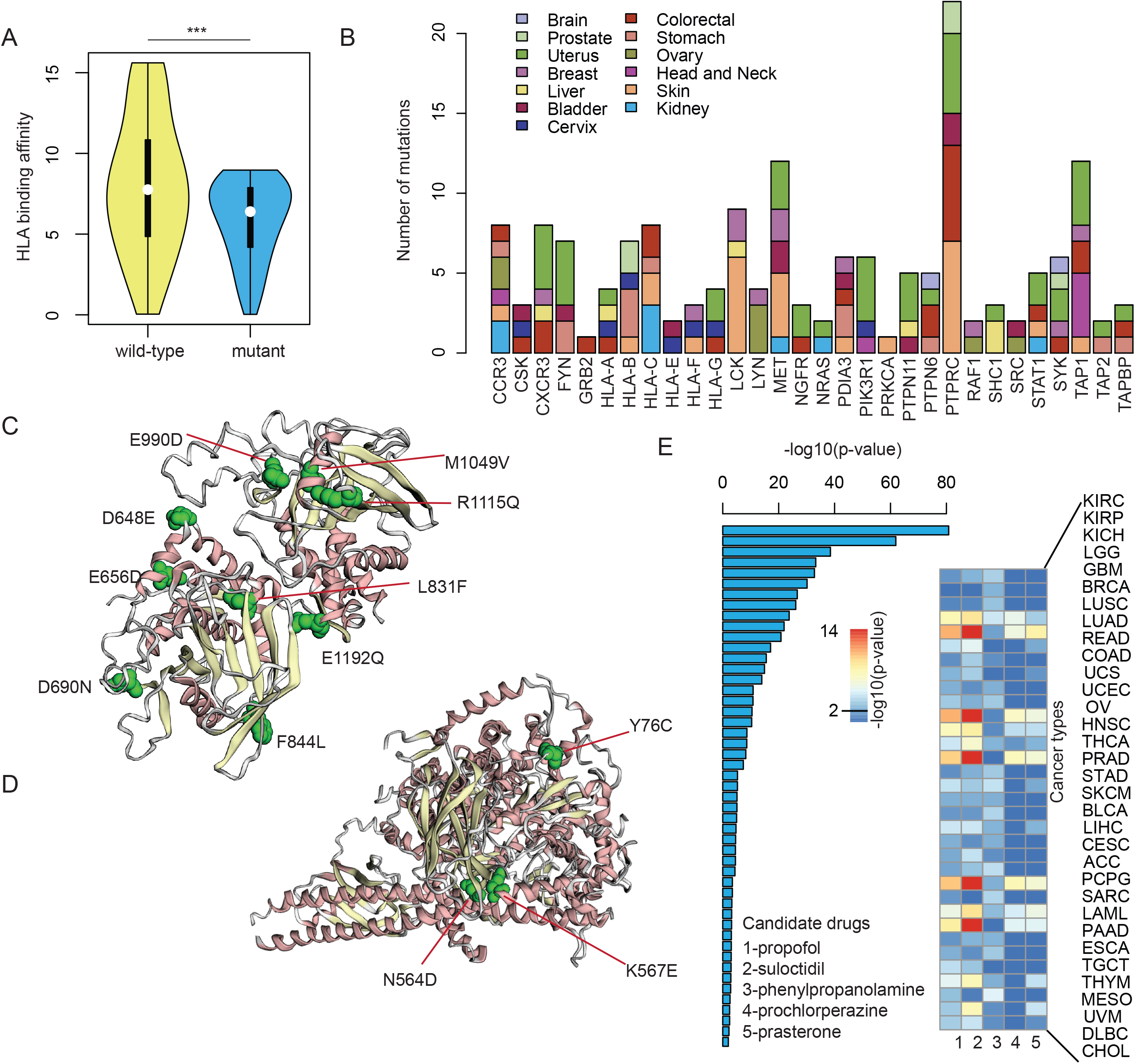
Candidate driver mutations and immune-response related small molecules. (A) The distribution of HLA binding affinity for wild-type (yellow) and mutated peptides (blue). ***p<0.001, Wilcox’s rank sum test. (B) The number of mutations that induced the decreased binding of HLAs across 30 genes. (C) The 3D structure of PTPRC and candidate mutations in this gene. (D) The 3D structure of PIK3R1 and candidate mutations in this gene. (E) Candidate immune-response related drugs and the corresponding p-values in cancer types predicted by network-based method.

Identification of candidate small molecule drugs is critical for the immunotherapy in cancer. Herein we identified candidate drugs that could increase the expression of identified immune gene signatures based on GSEA analysis (Figure 6A). We obtained more than 6,000 small molecules and their perturbation gene signatures from The Connectivity Map (71). We identified 49 small molecules that significantly perturbed the expression of immune gene signatures (p<0.05, Figure 7E). Interestingly, we identified propofol as the top one small molecule for increased immune response. Evidence had shown that propofol could possibly induce a favorable immune response in terms of preservation of IL-2/IL-4 and CD4+/CD8+ T cell ratio in the perioperative period for breast cancer (72). In vitro and in vivo studies had also suggested that propofol could independently reduce migration of cancer cells and metastasis (73). Taken together, these results suggest that the network-based integrative model not only can identify immune response-related gene signatures, it also helps prioritize candidate drugs for immunotherapy in cancer.

## Discussion

Systematically understanding the regulation of immune systems opens more avenues for cancer immunotherapy with a potent clinical efficacy. By integration of pan-cancer omics datasets, we have examined human tumor correlations in the context of immune-related pathways to an extent that is not previously possible. High-throughput gene expression data have been widely used to investigate differentially expressed genes in various types of cancer (74,75). Strikingly, we observed extensive immune-gene signatures with widespread perturbation in cancer compared with normal samples. These deregulated immune-related genes are enriched in ‘antigen processing and presentation’, ‘cytokines’ and ‘cytokines receptors’ pathways. Cytokines are critical molecular messengers for cells of the immune system to communicate with one another, which can directly stimulate immune effector cells and enhance tumor cell recognition. Increasing studies have demonstrated that cytokines have broad anti-tumor activity (76,77). Our analysis revealed that there are widespread perturbations of cytokine pathways in more than 80% of cancer types. A more detailed understanding of cytokine pathway regulation will provide new opportunities for improving cancer immunotherapy.

It is still unclear which factors are key players in regulating the expression of immune-related genes. Thus, we systematically analyzed two types of genetic regulation (including CNVs and eQTLs) in immune genes (Figure S6). We found that a number of immune-related genes recurrently dysregulated in various types of cancer. For example, up-regulation of BIRC5 is correlated with CNV amplification (78); and down-regulation of CCL14 is correlated with CNV deletion across cancer types. Moreover, eQTLs analysis revealed a hotspot locus that modulates the expression of DEFB1 (Figure S7), which might function through perturbation of CTCF binding. While a number of immune-related genes show correlation between copy number variations/mutations and gene expression, the relationship is weak and could not fully explain immune transcriptome perturbations. Thus, it is likely that other genetic (such as post-transcriptional regulation) or epigenetic variables (such as DNA methylation or histone modification) also contribute to the transcriptome perturbations in cancer.

Although several studies have proposed that the immune score, MHC score and CYT score are correlated with immune response, we found that these scores are complementary with each other. Thus, we proposed an integrated network model to predict the critical genes for immune response. This method identified several well-known immune response-related genes, such as B2M (79), TAP1 (80) and TAPBP (81), as well as several novel genes. HLA binding analysis also showed that these mutations are likely to induce the decreased binding of HLAs. Especially, we identified 165 candidate driver mutations that potentially play central roles in immune response. The mutations in PIK3R1 were validated in cell lines and were found to induce cancer cell growth. Moreover, we predicted candidate small molecules for immunotherapy based on whether the treatment could induce the expression of these signature immune genes. In particular, propofol was identified as the top-ranked drug candidates for immunotherapy. Propofol is the most commonly used in clinical anesthesia and propofol exhibits a good inhibitory effect on tumor recurrence and metastasis (82). Our current study found that propofol likely induces the expression of immune response-related genes, which may contribute to improved efficacy of immunotherapy.

Taken together, our integrative analysis of the human immunome across cancer types reveals several candidate gene and mutation markers that may be possible to use for predicting effective immune response prior to immunotherapy. Future studies will also need to evaluate the potential molecular functions of our predictive gene and mutation markers by low-throughput experiments.

## Authors’ Contributions

Conception and design: Y.L. and S.Y.

Development of methodology: Y.L., N.S. and S.Y.

Acquisition of data: Y.L., S.S, C.J.W., B.L. and S.Y.

Analysis and interpretation of data (e.g., statistical analysis, biostatistics, computational analysis): Y.L., B.L., N.S. and S.Y.

Writing, review, and/or revision of the manuscript: Y.L., T.T., D.J.M., E.W., A.C., S.G.E., G.G., B.L., N.S. and S.Y.

Administrative, technical, or material support (i.e., reporting or organizing data, constructing databases): T.T., B.B., M.S., D.Q., S.S., A.C.

Study supervision: S.Y.

## Acknowledgments

The authors would like to acknowledge Kevin Drew and Edward M. Marcotte from University of Texas at Austin for helpful discussions. The authors are grateful to contributions from TCGA Research Network Analysis Working Group, and also acknowledge the Biomedical Research Computing Facility at UT Austin and Texas Advanced Computing Center (TACC) for high-performance computing assistance.

